# The ITS region provides a reliable DNA barcode for identifying reishi/lingzhi (*Ganoderma*) from herbal supplements

**DOI:** 10.1101/2020.07.15.204073

**Authors:** Tess Gunnels, Matthew Creswell, Janis McFerrin, Justen B. Whittall

**Author notes:** Corresponding Author (JBW).

## Abstract

The dietary supplement industry is a growing enterprise, valued at over $100 billion by 2025 yet, a recent study revealed that up to 60% of herbal supplements may have substituted ingredients not listed on their labels, some with harmful contaminants. Substituted ingredients make rigorous quality control testing a necessary aspect in the production of supplements. Traditionally, species have been verified morphologically or biochemically, but this is not possible for all species if the identifying characteristics are lost in the processing of the material. One approach to validating plant and fungal ingredients in herbal supplements is through DNA barcoding complemented with a molecular phylogenetic analysis. This method provides an efficient, objective, rigorous and repeatable method for species identification. We employed a molecular phylogenetic analysis for species authentication of the commonly used fungal supplement, reishi (*Ganoderma lingzhi*), by amplifying and sequencing the nuclear ribosomal internal transcribed spacer regions (ITS) with genus-specific primers. PCR of six powdered samples and one dried sample sold as *G. lucidum* representing independent suppliers produced single, strong amplification products in the expected size-range for *Ganoderma*. Both best-hit BLAST and molecular phylogenetic analyses using a reference panel assembled from Genbank clearly identified the predominant fungal DNA was *G. lingzhi* in all seven herbal supplements. We detected variation in ITS among our samples, but all samples still fall within a large clade of *G. lingzhi*. ITS is a successful and cost-effective method for DNA-based species authentication that could be used in the herbal supplement industry for this and other fungal and plant species that are otherwise difficult to identify.

## Introduction

Barcoding is an efficient molecular tool for identifying morphologically, anatomically and biochemically enigmatic samples [1]. It has been applied across the tree of life [2, 3] and is increasingly employed in identifying the provenance of unidentifiable food products in restaurants [4] and in retail [5–7]. Leveraging the DNA content of processed living organisms that is not otherwise identifiable holds great prospects for quality control - especially helpful for maintaining the validity of active ingredients and avoiding contaminants that may cause allergic reactions for consumers [8]. This is particularly relevant in the herbal supplement industry, where safety and effectiveness are loosely regulated by the FDA through the Dietary Supplement Health and Education Act of 1994, which requires the manufacturer to ensure the safety and effectiveness of a supplement [9]. The Federal Food, Drug, and Cosmetic Act requires that manufacturers and distributors who wish to market dietary supplements that contain "new dietary ingredients" (not marketed in a dietary supplements before October 15, 1994) notify the Food and Drug Administration about these ingredients (see Section 413(d) of [10]). Under this Act, it is the responsibility of the manufacturer or distributor to assess whether a dietary supplement will be safe to use [10].

The herbal supplement industry is a growing enterprise, expected to amount to $104.78 billion dollars or more by 2025 [11, 12], yet, a recent study revealed that up to 60% of herbal supplements have substituted ingredients not listed on their labels, some of which are harmful contaminants [8]. For both marketing advantage and ethical concerns, suppliers must ensure accurate identification of all ingredients in their products [9]. Moreover, dietary supplement regulations require a manufacturer to perform identity testing on 100% of incoming lots of dietary ingredients, except when it has petitioned the FDA for a special exemption [10, 13]. For some manufactures, accurate identification of species, complete listing of ingredients, and precise reporting of potency are paramount. Furthermore, retailers are expected to exercise due diligence regarding oversight of suppliers. This is especially important since a large portion of the population consuming herbal supplements are doing so because their health is already compromised [14].

Reishi is one of the oldest herbal medicines in recorded history [reviewed in 15, 16] and estimated to represent 2% of the herbal supplement industry [14]. It is recommended as an anti-inflammatory and to enhance immunity [17, 18]. After being cultivated on rice, most reishi products are ground to a powder and sold in capsules as herbal supplements. Although the glossy, lignicolous, leathery, shelf-like polypore fruiting bodies of this group of laccate *Ganoderma* species are distinctive when fresh, once pulverized along with the rice medium (which often constitutes >50% of the dry weight), the powder is not easily differentiable morphologically, anatomically or biochemically (yet see [18, 19] for biochemical profiles of reishi and close relatives). Using biochemistry, Wu et al. [19] could only verify 26% of 19 reishi supplements purchased in the United States as true reishi (they use “*G. lucidum*”).

Adding to the difficulty of identifying processed reishi is the taxonomic confusion surrounding the genus *Ganoderma* [20]. The genus consists of approximately 80 species that fall into five or six clades - one of which is centered around *G. lucidum* sensu lato [21–26]. Because of their wood-decaying capabilities, several *Ganoderma* species have been investigated for biopulping [14] and bioremediation [27], however, it is most prized as a “model medicinal mushroom” [28] because of the putative health benefits of the triterpenoids and polysaccharides [29, 30]. The *G. lucidum* clade consists of several species that are in taxonomic flux [22, 24, 25]. According to a thorough morphological and molecular investigation of *G. lucidum* and *G. lingzhi*, “the most striking characteristics which differentiate *G. lingzhi* from *G. lucidum* are the presence of melanoid bands in the context, a yellow pore surface and thick dissepiments (80–120 μm) at maturity” [23]. *Ganoderma lucidum* sensu lato (including *G. ‘tsugae’*) can be found in the wild from Europe to northeastern China (likely escaped from cultivation in California and Utah, see [22]), yet *G. lingzhi* is restricted to Asia [18]. Fresh *G. lingzhi* has higher levels of triterpenoids which likely confer its physiological effects in humans [18, 19]. According to several recent molecular phylogenetic studies, the taxonomy of *G. lucidum* and *G. lingzhi* remains uncertain [18, 26, 31]. Most recent authors identify two distinct lineages which are supported by morphology [23], biochemistry [18], and molecular phylogenetics [22, 23, 24, 32].

The nuclear ribosomal ITS region is a powerful tool for barcoding plants and fungi [33–35]. It consists of two hypervariable spacers of approximately 200-250bp flanked by the 18S (small) subunit and 28S (large) subunit rDNA and separated by the 5.8S rDNA [36, 37]. Primers designed to bind to highly conserved portions of the 18S and 28S subunits have been widely used across plants and fungi [37]. However, lineage-specific primers have been developed for many groups of fungi in order to improve PCR success [23]. Lineage-specific primers improve PCR success especially when working with compromised DNA templates that may be degraded, contain inhibitors, or be composed of a mixture of species. *Ganoderma*-specific primers developed by Cao et al. [23] have been shown to improve PCR specificity. These primers have been used for barcoding reishi herbal supplements in previous studies (see [38] with limited sampling of reishi samples (n=4) and [14] with a broader retail sampling (n=14), but unclear how many unique suppliers were represented in the latter).

Although considerable attention has been given to the identification of the best barcoding loci and the development of unique and creative applications (recently reviewed in [34]), less explicit attention has been paid to the analysis of the data. The two main approaches for analysis of the DNA sequences arising from barcoding investigations are similarity-based measures (e.g., best-hit BLAST or nearest neighbor analysis) and phylogenetic methods (e.g., maximum likelihood or Bayesian tree-building algorithms). Some studies rely solely on best-hit BLAST [39] or otherwise crude phylogenetic approaches sometimes without assessment of the uncertainty (e.g., neighbor joining [40, 41]; or neighbor joining without bootstrap analysis [8, 42]). Similarity-based measures are known to fail in several common situations such as variable rates of molecular evolution [43, 44], gene duplication [44, 45] and changes in the gene’s composition [46]. Empirically, similarity-based approaches and phylogenetic methods for barcoding analysis are rarely compared explicitly even though they can produce conflicting identifications (see [47] which shows distance-based analyses performed best on Australian grasses, yet [48] who found tree-based methods worked better in *Vicia*).

Herein, we present an efficient barcoding method for unambiguous identification of the herbal supplement, reishi (*G. lingzhi*). We report successful DNA extraction, PCR amplification, and DNA sequencing of the ITS region from store-bought reishi samples. We compare the results from best-hit BLAST with two molecular phylogenetic approaches to determine if the species in the store-bought samples are correctly labeled or not.

## Materials and Methods

### Sampling

Store-bought samples were collected from multiple nutritional supplement retailers representing seven distinct suppliers of cultivated fungal products (Table 1). Four samples were encapsulated powders and two were loose powders, all of which purport to contain reishi, or *Ganoderma lucidum*, based on the product’s labeling. Of the seven supplements sampled, four were labeled as containing only mushroom mycelial biomass, two samples claimed to contain both mycelia and fruiting body, and one sample did not specify. The powdered samples varied in color, texture, and smell. All powdered samples are morphologically unidentifiable as a mushroom.

**Table 1.**
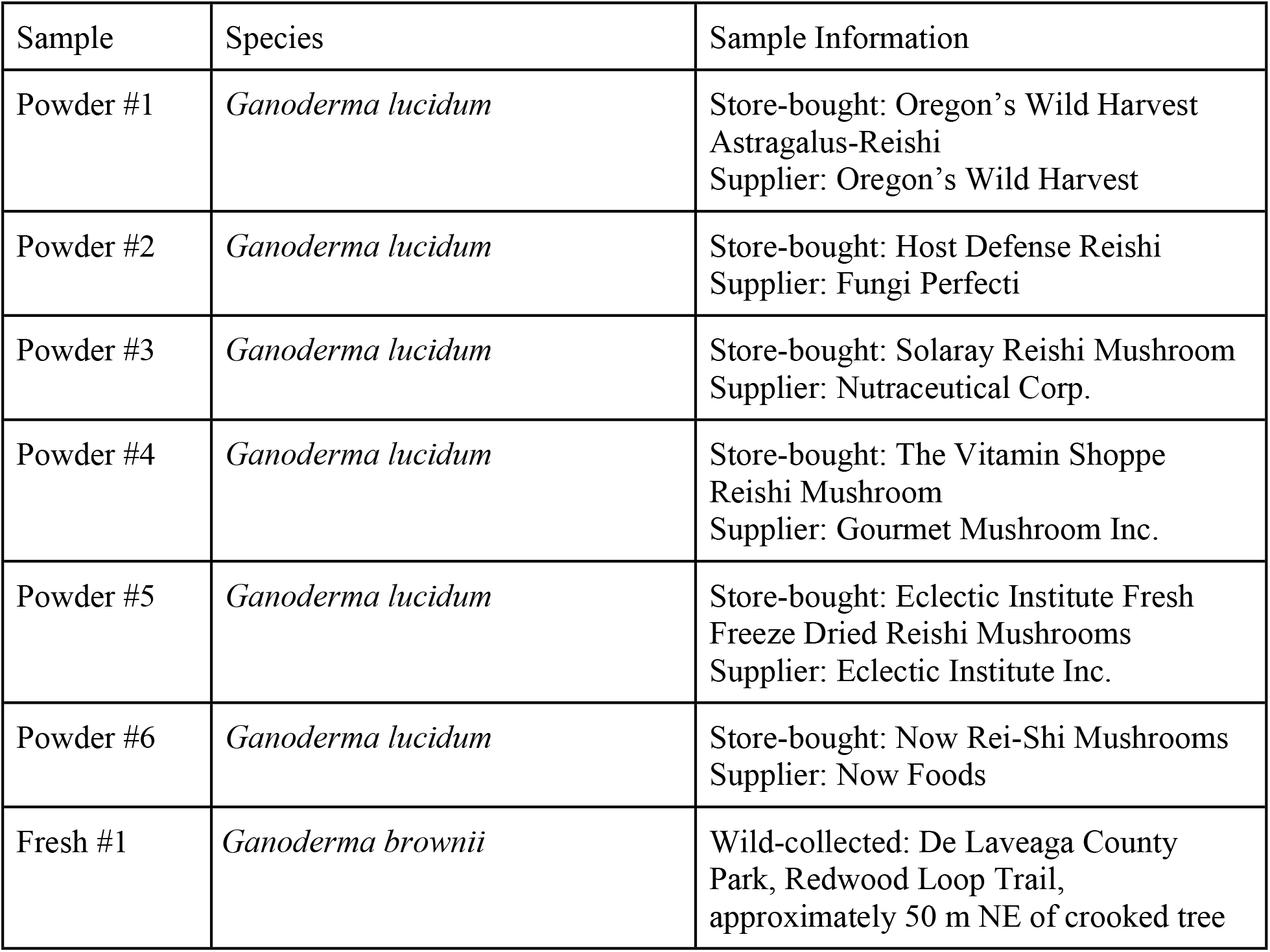
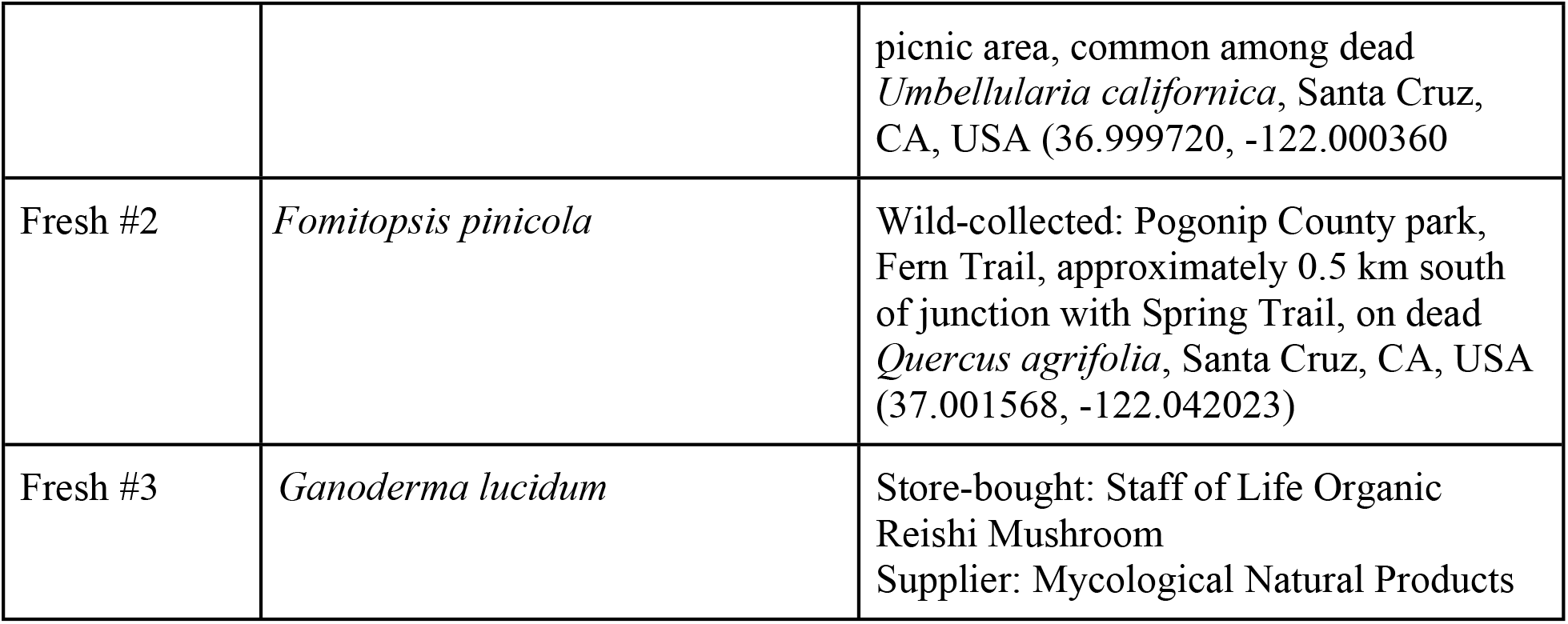
Sampling information for powdered and fresh samples that were store-bought or wild-collected.

| Sample | Species | Sample Information |
| --- | --- | --- |
| Powder #1 | *Ganoderma lucidum* | Store-bought: Oregon's Wild Harvest<br>Astragalus-Reishi<br>Supplier: Oregon's Wild Harvest |
| Powder #2 | *Ganoderma lucidum* | Store-bought: Host Defense Reishi<br>Supplier: Fungi Perfecti |
| Powder #3 | *Ganoderma lucidum* | Store-bought: Solaray Reishi Mushroom<br>Supplier: Nutraceutical Corp. |
| Powder #4 | *Ganoderma lucidum* | Store-bought: The Vitamin Shoppe<br>Reishi Mushroom<br>Supplier: Gourmet Mushroom Inc. |
| Powder #5 | *Ganoderma lucidum* | Store-bought: Eclectic Institute Fresh<br>Freeze Dried Reishi Mushrooms<br>Supplier: Eclectic Institute Inc. |
| Powder #6 | *Ganoderma lucidum* | Store-bought: Now Rei-Shi Mushrooms<br>Supplier: Now Foods |
| Fresh #1 | *Ganoderma brownii* | Wild-collected: De Laveaga County<br>Park, Redwood Loop Trail,<br>approximately 50 m NE of crooked tree |
|  |  | picnic area, common among dead *Umbellularia californica*, Santa Cruz, CA, USA (36.999720, -122.000360) |
| Fresh #2 | *Fomitopsis pinicola* | Wild-collected: Pogonip County park, Fern Trail, approximately 0.5 km south of junction with Spring Trail, on dead *Quercus agrifolia*, Santa Cruz, CA, USA (37.001568, -122.042023) |
| Fresh #3 | *Ganoderma lucidum* | Store-bought: Staff of Life Organic Reishi Mushroom<br>Supplier: Mycological Natural Products |

A fresh mushroom sample advertised as “organic reishi mushroom” was collected from the bulk herb section of Staff of Life natural goods store, Santa Cruz, CA in July of 2018 and was also evaluated based on its morphological characteristics (Fresh #3 in Table 1). The sample had been cut into strips of approximately 6 × 1 cm from cross sections of the fruiting body. The sample appeared woody in texture with extensive pore-containing regions similar to morphologically identified samples of the complete fruiting body.

Two additional fresh samples were collected from the wild (Santa Cruz County, CA, USA; Table 1) and used as positive controls for DNA extraction, PCR and sequencing (Table 1, Fresh #1 and Fresh #2). Samples were morphologically identified as *Ganoderma brownii* and *Fomitopsis pinicola* [49]. These two genera can be distinguished by the presence (*Ganoderma*) or absence (*Fomitopsis*) of bruising on the white pores of the fruiting body’s underside [49]. All samples were stored at room temperature until they were used for extraction.

### DNA extraction

Each nutritional supplement was extracted twice. Encapsulated samples were opened and only the powder contained within was used. Field collected samples were dissected and cut into smaller pieces for further morphological evaluation and then prepared for DNA extraction. Fresh tissue was removed from the underside of the fruiting body and cut into 2mm × 5mm rectangles for homogenization. Approximately 30-100 mg of material was homogenized in QIAGEN’s DNEasy Plant Mini Kit extraction buffer using a BeadBeater with 4 × 3.2 mm steel beads in XXTuff 2mL O-ring screw cap tubes (Biospec, Bartlesville, OK, USA). Following homogenization, DNA extraction was performed using the Qiagen DNEasy Plant Mini Kit following the manufacturer’s protocol (QIAGEN, Valencia, CA, USA). Concentration and purity of extracted DNA was evaluated using a Nanodrop spectrophotometer (NanoDrop Technologies Inc., Wilmington, DE, USA).

### PCR and sequencing

Several fungal ITS primer pairs were tested for initial success of amplification for both fresh and powdered samples (Table 2). Among them, the *Ganoderma*-specific primers (G-ITS-F1 and G-ITS-R2) were selected based on consistently producing strong single bands [23]. These primers were designed to prevent amplification from plant or other fungal DNA, which is a common problem with herbal supplements since they often include a plant-based growing medium.

**Table 2.**
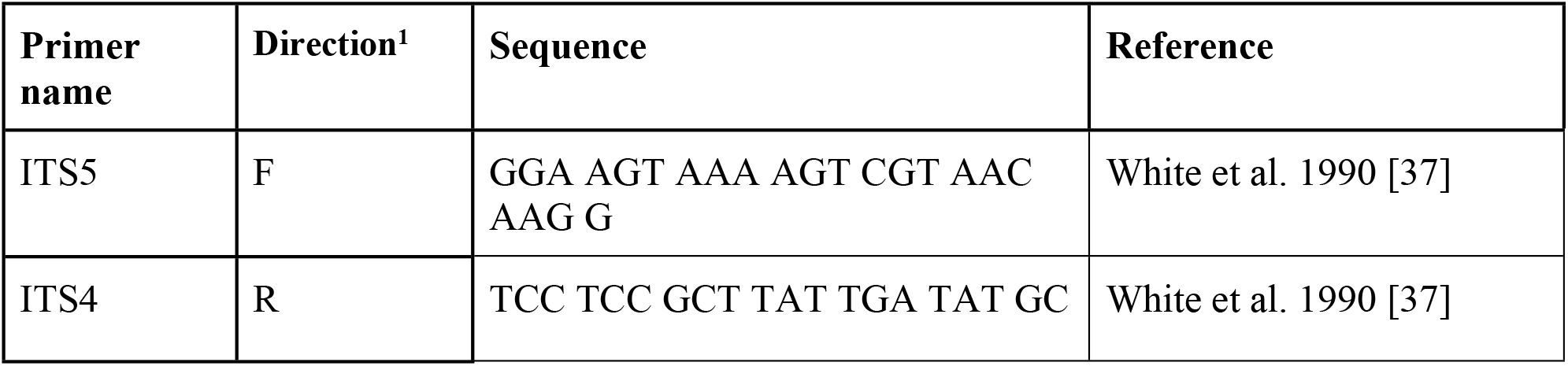
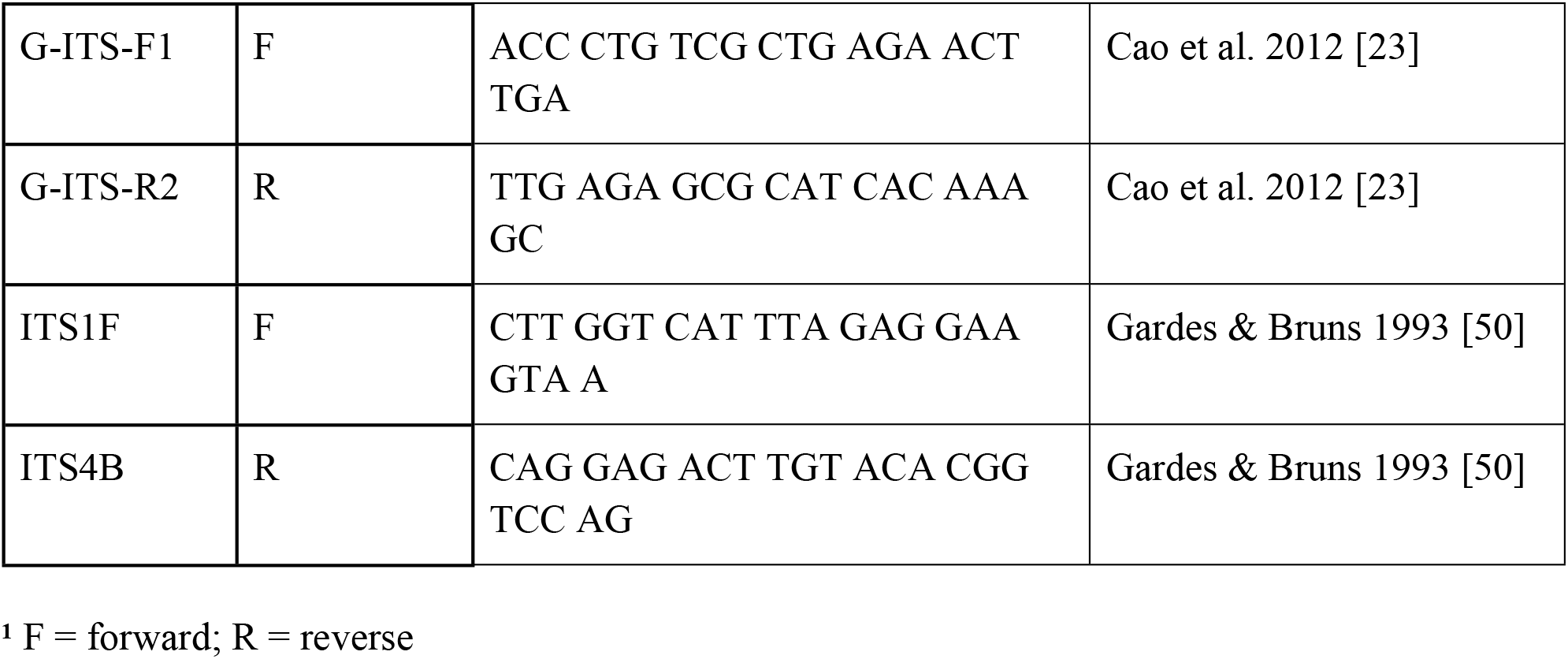
Three ITS primer pairs tested for amplification from reishi herbal supplements.

Extracted DNA was used as a template in 25 μL PCR reactions. Each reaction consisted of 2.5 μL of MgCl_2_ (25 mM), 2.5 μL of Taq Buffer B (Mg-free; 10X) (New England Biolabs, Ipswich, MA, USA), 2.5 μL of dNTPs (2.5 mM of each base), 2.5 μL of each of the aforementioned primers (10 μM), 0.25 μL of Taq polymerase (5U/μL) (New England Biolabs, Ipswich, MA, USA) and 1μL of extracted template DNA. A negative control (Milli-Q water in place of DNA template) was included in each PCR to ensure there was no contamination.

Amplification took place under the following thermal cycling conditions: initial denaturation at 92 °C for 2 min followed by 35 cycles of 94 °C for 1 min, 55 °C for 45 s, 72 °C for 45 s and a final extension step at 72 °C for 5 min. The PCR products were run on a 1% agarose gel stained with ethidium bromide alongside a 100 bp ladder (New England Biolabs, Ipswitch, MA, USA).

PCR reactions producing single, strong bands, were cleaned-up using shrimp alkaline phosphatase and directly sequenced in both directions using the Applied Biosystems 3730xl DNA Analyzer with the same primers used in PCR (Applied Biosystems, Waltham, Massachusetts, USA). Direct sequencing followed by BigDye Terminator or BigDye Primer methodologies per manufacturer recommendations (Sequetech, Mountain View, CA, USA).

Forward and reverse chromatograms for each sample were trimmed to remove primer sequence and low quality sequence. Reads were then aligned to form a single contiguous sequence using the pairwise alignment tool in Geneious Prime (Geneious Prime 2019.0.4, Biomatters, Auckland, NZ).

### Data analysis

#### BLAST

We used Basic Local Alignment Search Tool (BLAST) as the first method of identification for each sample. We used the megablast algorithm to search the nucleotide (nr/nt) collection to find the closest match to our sequences (BLASTDBv4) [51–53]. We compared the BLAST results from full-length sequence queries to the BLAST results of sequences trimmed to the portion of the alignment with maximum overlap with the reference database that were used in our phylogenetic approach (see below).

#### Multiple sequence alignment

We assembled an alignment of related sequences from Genbank. We started by including our top BLAST hits from the full sequence search query described above. If multiple Genbank accessions had equal coverage and identity as the top hit, we took at least one representative of each species which appeared. We also added all the unique Reishi samples (*G. lingzhi* and *G. lucidum*) of ITS using ENTREZ. Finally, we included representatives of as many *Ganoderma* species we could find using a filtered discontiguous megablast allowing us to limit ourselves to the most highly similar *Ganoderma* accessions, and increase our taxonomic coverage with a diverse and comprehensive reference set for nucleotide alignment and subsequent phylogenetic analysis. Several outgroup sequences were chosen which included other mushroom species belonging to the same order, *Polyporales*.

Sequences were aligned using the Geneious alignment tool (Biomatters, Auckland, New Zealand). All sequences were trimmed to approximately the same size producing an alignment of consistent length across the available ITS sequences of the Genbank reference set. Sequences with 100% nucleotide match to another sequence of the same species were removed so that only one representative sequence remained to simplify later phylogenetic analyses (Supplemental Table 3). If a sequence had a 100% match to a sequence belonging to a different species, both sequences were kept in the alignment to represent the additional taxonomic diversity. After all sequences were trimmed to approximately the same length we repeated the BLAST analysis of each sample to determine if sequence length affected the identity of the unknown samples.

##### Phylogenetic analyses

Maximum likelihood analysis was performed using the RAxML plug-in for Geneious (RAxML 8.2.11) [54]. We applied the GTR + CAT + I model of evolution and employed a rapid bootstrapping algorithm using 1,000 bootstrap replicates. Additionally, the MrBayes 3.2 plugin was used to build a Bayesian phylogenetic inference using Markov chain Monte Carlo (MCMC) algorithm (MrBayes 3.2.6) [55]. The GTR substitution model was used with a proportion invariable, remaining gamma rate variation model. Bayesian analysis used 1,500,000 Markov chains, which were sampled every 750 generations and after a burn-in length of 750,000 samples. The intention of our study is not to disentangle the taxonomic uncertainty regarding *G. lucidum* sensu lato and *G. lingzhi*. Therefore, throughout the results and discussion we have chosen to report the scientific names as they are reported in Genbank although some of these have been suggested to be mislabeled (see the Supplemental Table in [56]).

### Results

#### DNA extraction

The average concentration of DNA in the nine samples was 34.1 ng/uL (range 3.9 to 175.2; S1 Table). The average purity of the DNA measured as the 260/280 ratio was 1.34 (range 0.66 to 1.91; S1 Table).

#### PCR and sequencing

To assess successful amplification of the ITS region from newly extracted fungal DNA, PCR with three different primer pairs was performed and samples were visualized with gel electrophoresis. All primer pairs produced visible bands of expected size for the *ITS* region for both fresh and powdered samples. The *Ganoderma*-specific primers were chosen for all other analyses. Nucleotide sequences were recovered from the *ITS* region from all of the samples.

After trimming the sequences, lengths ranged from 780 base pairs to 895 base pairs with an average of 854 base pairs. Quality scores (HQ%) for full contiguous sequences of the forward and reverse directions ranged from 75.4% to 96.5% and averaged 91.2%.

#### Data analysis

##### BLAST

Our first approach for sample identification was to query Genbank for the top BLAST hit using the full length ITS sequence (Table 3). Of the seven store-bought samples, all yielded a top BLAST result that matched their labeled genus and species (*G. lucidum*). Top BLAST hits changed to *G. lingzhi* for all fresh samples when using the trimmed sequences from the 620 bp alignment as described in more detail below (Table 3).

**Table 3.** BLAST results using full length ITS sequences compared to ITS sequences trimmed to the GenBank reference panel alignment (620 bp) used in phylogenetic analysis.

|  |  |  | Full Length |  |  |  | Trimmed to Alignment Length |  |  |
| --- | --- | --- | --- | --- | --- | --- | --- | --- | --- |
| Sample Name | Presumed Species<sup>1</sup> | Genbank Accession | Sequence Length (bp) | Top BLAST Hit<sup>2</sup> | GenBank Query Coverage | GenBank Percent Similarity | Top BLAST Hit<sup>2</sup> | GenBank Query Coverage | GenBank Percent Similarity |
| Powder #1 | *G. lucidum* | XXXXXXX | 824 | *G. lucidum* (MF476200.1) | 100% | 100% | *G. lingzhi* (MH160076.1) | 100% | 100% |
| Powder #2 | *G. lucidum* | XXXXXXX | 739 | *G. lucidum* (MF476201.1) | 99% | 99% | *G. lingzhi* (MH160076.1) | 99% | 99% |
| Powder #3 | *G. lucidum* | XXXXXXX | 868 | *G. lucidum* (MF476200.1) | 100% | 100% | *G. lingzhi* (MH160076.1) | 100% | 100% |
| Powder #4 | *G. lucidum* | XXXXXXX | 868 | *G. lucidum* (MF476200.1) | 100% | 100% | *G. lingzhi* (MH160076.1) | 100% | 100% |
| Powder #5 | *G. lucidum* | XXXXXXX | 865 | *G. lucidum* (MF476200.1) | 100% | 100% | *G. lingzhi* (MH160076.1) | 100% | 100% |
| Powder #6 | *G. lucidum* | XXXXXXX | 844 | *G. lucidum* (MF476200.1) | 100% | 99% | *G. lingzhi* (MH160076.1) | 100% | 100% |
| Fresh #1 | *G. brownii* | XXXXXXX | 848 | *G. australe* (MK968731.1) | 100% | 97% | *G. brownii* (MG279159.1) | 100% | 100% |
| Fresh #2 | *Fomitopsis pinicola* | XXXXXXX | 780 | *F. pinicola* (EF530947.1) | 100% | 99% | *F. pinicola* (EF530947.1) | 100% | 99% |
| Fresh #3 | *G. lucidum* | XXXXXX | 868 | *G. lucidum*<br>(MF476200.1) | 100% | 99% | *G. lingzhi*<br>(MH160076.1) | 100% | 100% |

<sup>1</sup>Presumed species is based on product label for store-bought samples and morphological identification [49] for wild-collected samples.
<sup>2</sup>All top BLAST hits had an E value of 0.0.

#### Multiple sequence alignment

To further assess the identity of our store-bought and field-collected samples, they were aligned with a reference panel (S2 Table). After trimming the alignment to the length of the shortest sequence in the reference panel and temporarily removing identical sequences (S3 Table), we created a final alignment of 88 sequences measuring 620 base pairs long with 51.6% identity (including outgroups). Among these unique sequences, the average pairwise percent identity is 87.4%. Within the *G. lingzhi* clade, there were 91 variable sites (mean pairwise identity = 99.3%). The average genetic identity between our store-bought samples and the most similar Genbank accession was 99.8% (range 99.5 - 100%).

#### Phylogenetic analyses

The maximum likelihood analysis yielded a moderately resolved tree. Of the 85 distinct branches in the maximum likelihood tree, 31 branches (36%) were greater than 70%, a commonly used cut-off for 95% reliability (Fig 1) [57]. There is a moderately supported *G. lingzhi* clade containing nearly all of the samples labeled *G. lingzhi* and *G. lucidum* and all seven of the store-bought herbal supplement samples (bootstrap = 78%) (Fig 1). We also reconstructed a strongly supported clade containing the real *G. lucidum, G. tsugae, G. oregonense* and *G. carnosum* (Clade B) (Fig. 1). We have applied clade names A and B from Loyd et al. [22] and Zhou et al. [24]. Clade A containing *G. tuberosum* and *G. multipileum* appears paraphyletic in Fig. 1, however the deepest nodes are very weakly supported (<10% bootstrap) and therefore, not in conflict with previous studies [22, 24].

**Fig 1.**
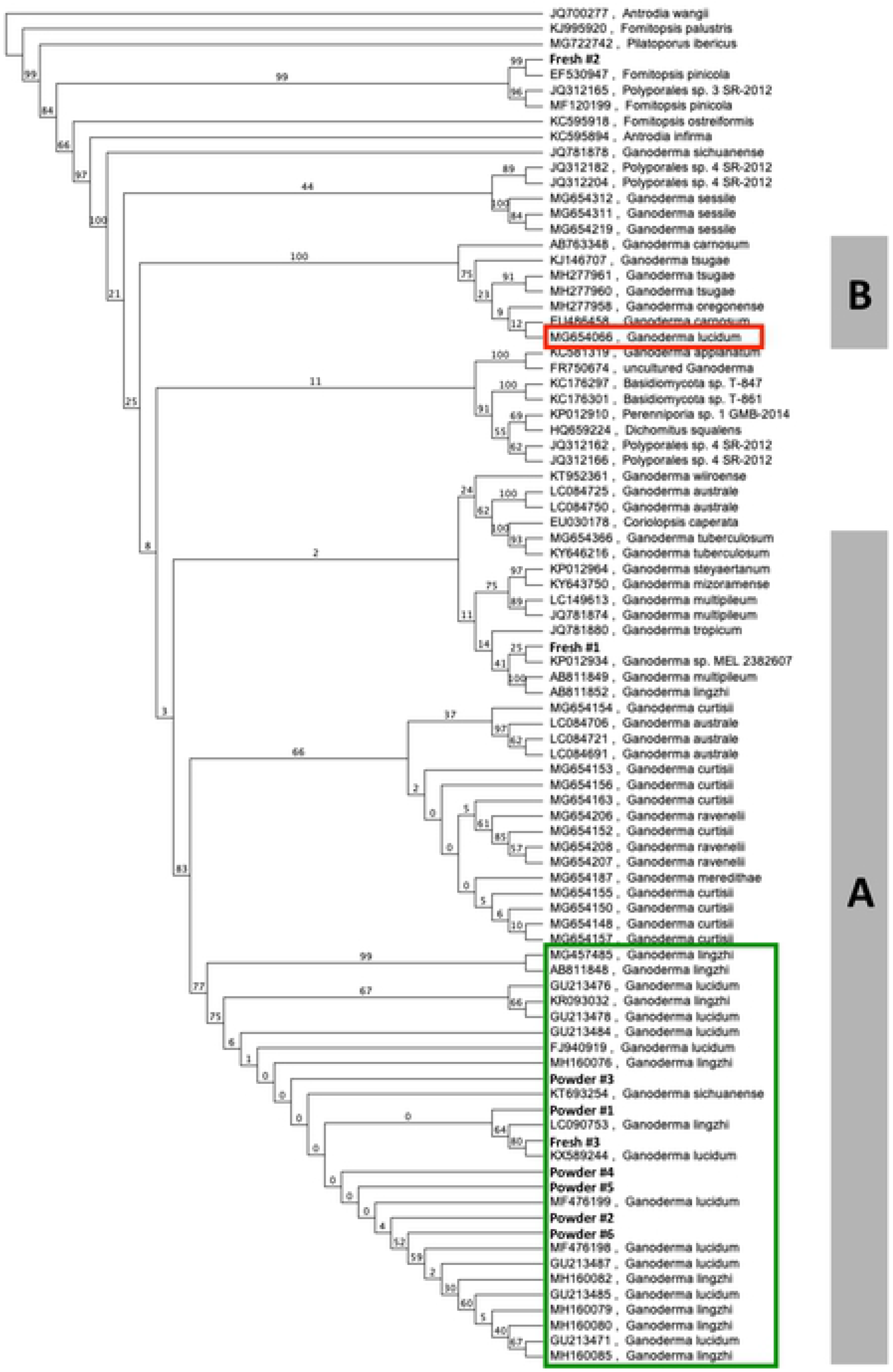
Maximum likelihood phylogenetic analysis. RAxML phylogeny including store-bought samples, wild collected samples and the Genbank reference set. Bootstrap values are shown above the branches. Identical sequences that were collapsed to a single representative are listed on the right. Clade names A and B are from Loyd et al. [22] and Zhou et al. [24]. The red rectangle identifies the only true *G. lucidum* sample per Loyd et al. [14] and the green rectangle contains the samples referred to as the *G. lingzhi* clade.

We detected ITS sequence variation among the store-bought reishi samples. Six of our seven store-bought samples (Powders #1-5 and Fresh #3) were phylogenetically allied with one of the two large clades of largely unresolved *G. lingzhi* samples in Clade A with very low (or no) bootstrap support therein (Fig 1). Among those six store-bought samples, only Fresh #3 had a sister relationship (to “*G. lucidum*” KX589244) with moderate bootstrap support (79%). The seventh store-bought sample, Powder #6, is sister to the other large clade of eight poorly resolved accessions named *G. lingzhi* or *G. lucidum* with weak support (bootstrap = 60%).

As a control, we included two fresh samples of wild-collected polypores. Fresh #1 was morphologically identified as *Ganoderma brownii* and falls clearly outside the *G. lingzhi* clade in a poorly resolved cluster of *Ganoderma* accessions in Clade A. Fresh #2 was morphologically identified as *Fomitopsis pinicola* and it strongly paired with a *F. pinicola* Genbank accession with 99% bootstrap support (EF530947; Fig 1).

The maximum likelihood tree revealed three phylogenetically aberrant Genbank accessions worth noting (Fig 1): (1) a *G. lingzhi* sample (AB811852) that is strongly supported as sister to *G. multipileum* (100% bootstrap) and deeply nested in Clade A; (2) a *G. lucidum* sample (MG654066) falls within a small, yet moderately supported clade of mostly North American samples - *G. tsugae*, *G. carnosum* (UK) and *G. oregonense* (bootstrap = 100%); (3) a *G. sichuanense* sample (KT693254) is nested within the moderately supported *G. lingzhi* clade (78% bootstrap; see [22] for discussion about this taxon).

The Bayesian phylogenetic analysis is largely consistent with the maximum likelihood tree, yet slightly less resolved. Of the 85 branches in the Bayesian tree, 23 of them (27%) are strongly supported with posterior probabilities greater than 0.95 (S1 Fig). There is a strongly supported clade containing nearly all of the Genbank accessions named *G. lucidum* and *G. lingzhi* and all of the store-bought samples (posterior probability = 0.987), yet no clear differentiation between the *G. lucidum* and *G. lingzhi* samples as named in Genbank (S1 Fig). Clade B is strongly supported as monophyletic (posterior probability = 0.9987) containing the only correctly identified *G. lucidum* accession (per [22]). The three putatively misidentified accessions described for the maximum likelihood analysis had similarly aberrant phylogenetic affinities in the Bayesian analysis (S1 Fig).

In the Bayesian phylogenetic tree, six of the seven store-bought samples are part of a large unresolved polytomy of accessions named *G. lucidum* and *G. lingzhi* (the true *G. lingzhi* clade). The exception, Fresh #3, falls within a subclade that is composed of an unresolved trichotomy with two other Genbank samples - one labeled *G. lucidum* and one labeled *G. lingzhi* (KX589244 and LC090753, respectively). Together, the three are supported as monophyletic with 0.9744 posterior probability (S1 Fig).

For the Bayesian analysis, the two control samples allied with similar Genbank accessions as in the maximum likelihood analysis. Fresh #1 (morphologically identified as *G. brownii*) falls clearly outside the true *G. lingzhi* clade in a highly unresolved cluster of *Ganoderma* species (S1 Fig). Fresh #2 (morphologically identified as *Fomitopsis pinicola)* is strongly supported as sister with a Genbank *F. pinicola* sample (EF530947; posterior probability = 0.9997; S1 Fig).

## Discussion

Our study demonstrates that the ITS region provides an efficient barcode for store-bought reishi herbal supplements as previously described by Loyd et al. [14] and Raja et al. [38]. Amplifiable genomic DNA was successfully extracted from both powdered and fresh samples - all of which were correctly identified. Loyd et al. [14] found widespread label confusion in both “grow your own” kits (15/17) and manufactured herbal supplements (13/14) that were sold as *G. lucidum*. They used both ITS and tef1-alpha sequences to identify the manufactured supplements were all *G. lingzhi*, except one *G. applanatum* [14]. The label confusion surrounding *G. lucidum* and *G. lingzhi* was likely unintentional due to the taxonomic uncertainty, although there are clear biochemical differences (and therefore potential human physiological consequences) that differentiate the two taxa) [18, 19]. In fact, Wu et al. [19] considered 26% of their 19 samples “verified” even though the labels read “*G. lucidum*” and not the correct species name, “*G. lingzhi*.” In general, herbal supplements are notoriously mislabeled - Newmaster et al.’s [8] study of plant herbal supplements found 59% (30 out of 44) had species substitutions and about 33% of these products had fillers or contaminants that were not listed on the product label - some of which could pose health risks to consumers. DNA barcoding will continue to be a valuable tool for manufacturers, retailers and consumer-watch groups, especially for herbal supplements like reishi where a lack of morphological and chemical distinctiveness once in powder form coupled with taxonomic confusion permeates the industry.

All of the samples we examined had BLAST and phylogenetic results suggesting they were clearly members of Clade A sensu Zhou et al. [24] and Loyd et al. [14] which only includes “*G. lucidum*” as defined in the broadest sense (see [22, 32] for what “*G. lucidum* sensu lato” refers to). None of our nine distinct distributors sampled contained material belonging to Clade B (*G. lucidum* sensu stricto). Technically all of our samples are misidentified since they are being sold as “*G. lucidum*”, yet are molecularly allied with the *G. lingzhi* samples in Clade A. A similar case of mistaken identity is reported by Loyd et al. [14]. We assume the mislabeling was unintentional and arose from the taxonomic confusion rife in this genus (yet recently and lucidly clarified by [32]). This level of mistaken identity (100%) is relatively rare among herbal supplement barcoding studies in general [8]. Although we only included seven store-bought samples, these represent seven distinct suppliers thereby broadening the implications of our study to all the retailers using those suppliers as well, something previous *Ganoderma* retail barcoding studies have not reported (using different retail samples from the same supplier could be considered pseudoreplication; see [14, 38]). Misidentifications can arise at any of the multitude of links that connect the growers with the retailers. Our targeted sampling at the supplier stage clearly indicates that the misidentifications are likely applied early in the process and inherited by the retailers.

Our study does not attempt to resolve the taxonomic ambiguity among closely related species within the genus *Ganoderma* which permeates the available sequence data in Genbank [23, 32, 56]. However, we do not wish to contribute to the confusion so will attempt to reconcile some of the Genbank names with the recent literature. Most importantly for reishi identification, we have recovered a very strongly supported *G. lucidum* sensu stricto (Clade B, bootstrap = 100%) and a more diffuse Clade A that includes the samples that should be identified as *G. lingzhi* according to Zhou et al. [24], Patterson & Lima [32], and Loyd et al. [22]. Clade A represents the medicinally important reishi (also known by the common name “lingzhi”) which is properly named *G. lingzhi* and restricted to Asia (see [32] for nomenclatural justification; also see Correction in [56]). Genbank sample MG654066 named *G. lucidum* (in Clade B) is the only sample that represents *G. lucidum* sensu stricto, native to Europe, closely related to North American *G. oregonense* and *G. tsugae*, and most likely introduced to Utah and California, USA according to Loyd et al. [22]. In comparison to Cao et al. [23] who examined four nuclear genes including ITS (yet only four samples of *G. lingzhi* and *G. lucidum* and a total of 13 species), our analysis has 10 *G. lingzhi* and 11 *G. lucidum* sensu lato samples) and more species overall (n = 29), yet limited to the single barcoding locus, the ITS region.

More broadly, our phylogenetic results (Fig 1) are generally congruent with previous studies employing the ITS region [22, 23, 24, 56]. They all report similar clades that we have identified in our results, yet often with higher confidence when including more loci. Although we have chosen to report the Genbank organism fields as they are in the database, we highlight the taxonomic confusion around these lineages and anticipate their realignment in Genbank in the near future.

ITS variation within the *G. lingzhi* clade allowed us to further partition our store-bought samples. Most samples were part of a large unresolved polytomy, but in two cases (one within the polytomy and one in the sister clade), there were distinct phylogenetic affinities clearly indicating separate sources. The intraspecific variation in ITS could prove valuable for tracing the intraspecific provenance of some reishi herbal supplements, but will likely need to be complemented with additional rapidly evolving loci (e.g., tef1-alpha, see [14, 22, 24]).

Methodologically, BLAST and phylogenetic analyses agreed on the provenance of all of the store-bought samples. When the rates of molecular evolution are relatively constant among the samples, in the absence of gene duplication, and when gene structure is conserved (such as for the ITS region), BLAST and phylogenetic methods are predicted to converge on similar identifications [43, 44, 46]. However, when any of those characteristics are violated, similarity based approaches, such as BLAST, that rely on a local alignment algorithm (some modification of [58]) can be misleading. Alternatively, phylogenetic analysis relies on a global alignment algorithm [59] spanning the entire length of the locus being compared and is more likely to identify the evolutionary history of the samples for that locus [44], yet is most rigorously employed with a model-based approach (e.g. maximum likelihood and bayesian methodologies) compared to a distance-based approach that is commonly found in the barcoding literature, yet fraught with weaknesses. Our results were generally robust to whether we used the entire ITS region or the trimmed region of overlap used in the multiple sequence alignment and subsequent phylogenetic analysis (all results point to *G. lingzhi* in Clade A). However, because of the nomenclatural issue associated with many Genbank samples, it appears that our results change from *G. lucidum* to *G. lingzhi*. This points to the importance of rectifying the Genbank taxonomy to avoid future, honest misidentifications.

## Acknowledgements

The authors are grateful to the lab support staff in the Department of Biology that provided essential services throughout this study. Oregon Wild Harvest (Redmond, OR) supported TG throughout the duration of the study. Jonathan Eisen (UC Davis) kindly helped with references to the BLAST vs. phylogenetic analysis discussion.

## Supporting Information

**S1 Fig. Bayesian phylogenetic analysis.** Mr. Bayes tree with posterior probabilities indicated along the branches. Genbank Accession numbers are indicated for reference panel. Samples are identified with reference to Table 1.

**S1 Table. DNA concentration and purity for herbal supplement powder samples and fresh samples.**

**S2 Table. Genbank reference panel sampling.**

## References

1. Hebert PD, Cywinska A, Ball SL, Dewaard JR. Biological identifications through DNA barcodes. Proc Biol Sci. 2003 Feb 7;270(1512):313–21.

2. The iBol Consortium. 2020 [cited 2020 June 17]. iBol [Internet]. Available from: https://ibol.org/about/ibol-consortium/

3. International Barcode of Life. 2020 [cited 2020 June 17]. iBol [Internet]. Available from: https://ibol.org/about/our-vision/

4. Christiansen H, Fournier N, Hellemans B, Volckaert FA. Seafood substitution and mislabeling in Brussels’ restaurants and canteens. Food control. 2018 Mar 1;85:66–75.

5. Hu Y, Huang SY, Hanner R, Levin J, Lu X. Study of fish products in Metro Vancouver using DNA barcoding methods reveals fraudulent labeling. Food Control. 2018 Dec 1;94:38–47.

6. Pardo MÁ, Jiménez E, Viðarsson JR, Ólafsson K, Ólafsdóttir G, Daníelsdóttir AK, Pérez-Villareal B. DNA barcoding revealing mislabeling of seafood in European mass caterings. Food Control. 2018 Oct 1;92:7–16.

7. Luque GM, Donlan CJ. The characterization of seafood mislabeling: A global meta-analysis. Biol Conserv. 2019 Jun 21.

8. Newmaster SG, Grguric M, Shanmughanandhan D, Ramalingam S, Ragupathy S. DNA barcoding detects contamination and substitution in North American herbal products. BMC Med. 2013 Dec 1;11(1):222.

9. Dodge T, Litt D, Kaufman A. Influence of the dietary supplement health and education act on consumer beliefs about the safety and effectiveness of dietary supplements. J Health Commun. 2011 Feb 28;16(3):230–44.

10. New dietary ingredients in dietary supplements-background for industry. 2019 Oct 4 [cited 2020 June 17]. US FDA [Internet]. Available from: https://www.fda.gov/food/new-dietary-ingredients-ndi-notification-process/new-dietary-ingredients-dietary-supplements-background-industry

11. Press release: herbal supplements market analysis, size, share, growth, trends and forecast to 2025. 2020 June 10 [cited 2020 June 17]. In: Market Watch [Internet]. Available from: https://www.marketwatch.com/press-release/global-herbal-supplements-market-size-growth-status-and-forecast-2020-2025-2020-06-10

12. Binns CW, Lee MK, Lee AH. Problems and prospects: public health regulation of dietary supplements. Annu Rev Public Health. 2018 Apr 1;39:403–20.

13. Electronic Code of Federal Regulations (e-CFR) Title 21, Chapter I, Subchapter B, Part 111. 2020 June 15. [cited 2020 June 17]. Available from: https://www.ecfr.gov/cgi-bin/text-idx?SID=9118251184f152d3b6f8bc44968863eb&mc=true&node=pt21.2.111&rgn=div5#sp21.2.111.a

14. Loyd AL, Richter BS, Jusino MA, Truong C, Smith ME, Blanchette RA, Smith JA. Identifying the “mushroom of immortality”: assessing the *Ganoderma* species composition in commercial Reishi products. Frontiers in Microbiology 2018 July 16; 9:1557.

15. Mizuno T, Wang G, Zhang J, Kawagishi H, Nishitoba T, Li J. Reishi, *Ganoderma lucidum* and *Ganoderma tsugae*: bioactive substances and medicinal effects. Food Rev Int. 1995 Feb 1;11(1):151–66.

16. Sliva D. Cellular and physiological effects of *Ganoderma lucidum* (Reishi). Mini Rev Med Chem. 2004 Oct 1;4(8):873–9.

17. Wang XC, Xi RJ, Li Y, Wang DM, Yao YJ. The species identity of the widely cultivated *Ganoderma*,‘*G. lucidum’*(Ling-zhi), in China. PLoS One. 2012;7(7).

18. Hennicke F, Cheikh-Ali Z, Liebisch T, Maciá-Vicente JG, Bode HB, Piepenbring M. Distinguishing commercially grown *Ganoderma lucidum* from *Ganoderma lingzhi* from Europe and East Asia on the basis of morphology, molecular phylogeny, and triterpenic acid profiles. Phytochemistry. 2016 Jul 1;127:29–37.

19. Wu DT, Deng Y, Chen LX, Zhao J, Bzhelyansky A, Li SP. Evaluation on quality consistency of *Ganoderma lucidum* dietary supplements collected in the United States. Sci Rep. 2017 Aug 10;7(1):1–0.

20. Moncalvo JM, Wang HH, Hseu RS. Phylogenetic relationships in *Ganoderma* inferred from the internal transcribed spacers and 25S ribosomal DNA sequences. Mycologia. 1995 Mar 1;87(2):223–38.

21. Hibbett DS, Donoghue MJ. Progress toward a phylogenetic classification of the Polyporaceae through parsimony analysis of mitochondrial ribosomal DNA sequences. Can J Bot. 1995 Dec 31;73(S1):853–61.

22. Loyd AL, Barnes CW, Held BW, Schink MJ, Smith ME, Smith JA, Blanchette RA. Elucidating" lucidum": Distinguishing the diverse laccate *Ganoderma* species of the United States. PloS One. 2018;13(7).

23. Cao Y, Wu SH, Dai YC. Species clarification of the prize medicinal *Ganoderma* mushroom “Lingzhi”. Fungal Divers. 2012 Sep 1;56(1):49–62.

24. Zhou LW, Cao Y, Wu SH, Vlasák J, Li DW, Li MJ, Dai YC. Global diversity of the *Ganoderma lucidum* complex (Ganodermataceae, Polyporales) inferred from morphology and multilocus phylogeny. Phytochemistry. 2015 Jun 1;114:7–15.

25. Dai YC, Zhou LW, Hattori T, Cao Y, Stalpers JA, Ryvarden L, Buchanan P, Oberwinkler F, Hallenberg N, Liu PG, Wu SH. *Ganoderma lingzhi* (Polyporales, Basidiomycota): the scientific binomial for the widely cultivated medicinal fungus Lingzhi. Mycol Prog. 2017 Dec 1;16(11-12):1051–5.

26. Liao B, Chen X, Han J, Dan Y, Wang L, Jiao W, Song J, Chen S. Identification of commercial *Ganoderma (Lingzhi)* species by ITS2 sequences. Chin Med. 2015 Dec 1;10(1):22.

27. Matos AJ, Bezerra RM, Dias AA. Screening of fungal isolates and properties of *Ganoderma applanatum* intended for olive mill wastewater decolourization and dephenolization. Lett App Microbiol. 2007 Sep;45(3):270–5.

28. Chen S, Xu J, Liu C, Zhu Y, Nelson DR, Zhou S, Li C, Wang L, Guo X, Sun Y, Luo H. Genome sequence of the model medicinal mushroom *Ganoderma lucidum*. Nat Commun. 2012 Jun 26;3(1):1–9.

29. Jin X, Ruiz Beguerie J, Sze DM, Chan GC. *Ganoderma lucidum* (Reishi mushroom) for cancer treatment. Cochrane Database Syst Rev. 2012 Jun 13;(6):CD007731. Update in: Cochrane Database Syst Rev. 2016;4:CD007731.

30. Wachtel-Galor S, Yuen J, Buswell JA, Benzie IF. *Ganoderma lucidum* (Lingzhi or Reishi). InHerbal Medicine: Biomolecular and Clinical Aspects. 2nd edition 2011. CRC Press/Taylor & Francis.

31. Hapuarachchi KK, Wen TC, Deng CY, Kang JC, Hyde KD. Mycosphere essays 1: Taxonomic confusion in the *Ganoderma lucidum* species complex. Mycosphere. 2015 Jan 1;6(5):542–59.

32. Paterson RR, Lima N. Failed PCR of *Ganoderma* type specimens affects nomenclature. Phytochemistry. 2015 Jun 1;114:16–7.

33. Schoch CL, Seifert KA, Huhndorf S, Robert V, Spouge JL, Levesque CA, Chen W, Fungal Barcoding Consortium. Nuclear ribosomal internal transcribed spacer (ITS) region as a universal DNA barcode marker for *Fungi*. Proc Natl Acad Sci USA. 2012 Apr 17;109(16):6241–6.

34. Xu J. Fungal DNA barcoding. Genome. 2016;59(11):913–32.

35. Li DZ, Gao LM, Li HT, Wang H, Ge XJ, Liu JQ, Chen ZD, Zhou SL, Chen SL, Yang JB, Fu CX. (2011). Comparative analysis of a large dataset indicates that internal transcribed spacer (ITS) should be incorporated into the core barcode for seed plants. Proc Natl Acad Sci USA.;108:19641–6.

36. Baldwin BG. Phylogenetic utility of the internal transcribed spacers of nuclear ribosomal DNA in plants: an example from the Compositae. Mol Phylogenet Evol. 1992 Mar 1;1(1):3–16.

37. White TJ, Bruns T, Lee SJ, Taylor J. Amplification and direct sequencing of fungal ribosomal RNA genes for phylogenetics. PCR protocols: a guide to methods and applications. 1990 Jan 1;18(1):315–22.

38. Raja HA, Baker TR, Little JG, Oberlies NH. DNA barcoding for identification of consumer-relevant mushrooms: A partial solution for product certification?. Food Chem. 2017 Jan 1;214:383–92.

39. Günther B, Raupach MJ, Knebelsberger T. Full-length and mini-length DNA barcoding for the identification of seafood commercially traded in Germany. Food Control. 2017 Mar 1;73:922–9.

40. Pereira LH, Hanner R, Foresti F, Oliveira C. Can DNA barcoding accurately discriminate megadiverse Neotropical freshwater fish fauna?. BMC Genet. 2013 Dec 1;14(1):20.

41. Steinke D, Zemlak TS, Hebert PD. Barcoding Nemo: DNA-based identifications for the ornamental fish trade. PLoS One. 2009; 4(7).

42. Sultana S, Ali ME, Hossain MM, Naquiah N, Zaidul IS. Universal mini COI barcode for the identification of fish species in processed products. Food Res Int. 2018 Mar 1;105:19–28.

43. Eisen JA. Phylogenomics: improving functional predictions for uncharacterized genes by evolutionary analysis. Genome Res. 1998 Mar 1;8(3):163–7.

44. Koski LB, Golding GB. The closest BLAST hit is often not the nearest neighbor. J Mol Evol. 2001 Jun 1;52(6):540–2.

45. Eisen JA, Kaiser D, Myers RM. Gastrogenomic delights: a movable feast. Nat Med. 1997 Oct;3(10):1076.

46. Spouge JL, Mariño-Ramírez L. The practical evaluation of DNA barcode efficacy. In: DNA Barcodes. Totowa, NJ: Humana Press; 2012. pp. 365–377.

47. Birch JL, Walsh NG, Cantrill DJ, Holmes GD, Murphy DJ. Testing efficacy of distance and tree-based methods for DNA barcoding of grasses (Poaceae tribe *Poeae*) in Australia. PloS One. 2017;12(10).

48. Wu FF, Gao Q, Liu F, Wang Z, Wang JL, Wang XG. DNA barcoding evaluation of *Vicia* (Fabaceae): Comparative efficacy of six universal barcode loci on abundant species. J Syst Evol. 2020 Jan;58(1):77–88.

49. Arora D, Hershey H. Mushrooms demystified. Berkeley: Ten Speed Press; 1986.

50. Gardes M, Bruns TD. ITS primers with enhanced specificity for basidiomycetes‐application to the identification of mycorrhizae and rusts. Mol Ecol. 1993 Apr;2(2):113–8.

51. Altschul SF, Gish W, Miller W, Myers EW, Lipman DJ. Basic local alignment search tool. J Mol Biol. 1990 Oct 5;215(3):403–10.

52. Zhang Z, Schwartz S, Wagner L, Miller W. A greedy algorithm for aligning DNA sequences. J Comput Biol. 2000 Feb 1;7(1-2):203–14.

53. Morgulis A, Coulouris G, Raytselis Y, Madden TL, Agarwala R, Schäffer AA. Database indexing for production MegaBLAST searches. Bioinformatics. 2008 Aug 15;24(16):1757–64.

54. Stamatakis A. RAxML version 8: a tool for phylogenetic analysis and post-analysis of large phylogenies. Bioinformatics. 2014 May 1;30(9):1312–3.

55. Huelsenbeck JP, Ronquist F. MRBAYES: Bayesian inference of phylogenetic trees. Bioinformatics. 2001 Aug 1;17(8):754–5.

56. Jargalmaa S, Eimes JA, Park MS, Park JY, Oh SY, Lim YW. Taxonomic evaluation of selected *Ganoderma* species and database sequence validation. PeerJ. 2017 Jul 27;5:e3596.

57. Hillis DM, Bull JJ. An empirical test of bootstrapping as a method for assessing confidence in phylogenetic analysis. Syst Biol. 1993 Jun 1;42(2):182–92.

58. Smith TF, Waterman MS. Identification of common molecular subsequences. J Mol Biol. 1981 Mar 25;147(1):195–7.

59. Needleman SB, Wunsch CD. A general method applicable to the search for similarities in the amino acid sequence of two proteins. J Mol Biol. 1970 Mar 28;48(3):443–53.

